# DOTSeq enables genome-wide detection of differential ORF usage

**DOI:** 10.1101/2025.09.24.678201

**Authors:** Chun Shen Lim, Gabrielle S.W. Chieng

## Abstract

Protein synthesis is regulated by multiple *cis*-regulatory elements, including small ORFs, yet current differential translation methods assume uniform changes at the gene level. We present `DOTSeq`, a Differential ORF Translation statistical framework that resolves ORF-level regulation in bulk ribosome profiling (Ribo-seq) experiments and provides ORF-level read summarisation for single-cell Ribo-seq. `DOTSeq`’s core module, Differential ORF Usage (`DOU`), quantifies changes in an ORF’s relative contribution to a gene’s translation output, using a beta-binomial GLM with flexible dispersion modelling. `DOTSeq` also implements ORF-level Differential Translation Efficiency (`DTE`) using a standard approach to complement `DOU`. Benchmarks show that `DOU` achieves superior sensitivity with near-nominal FDR across effect sizes, while `DTE` and some existing methods excel when technical noise is low. `DOTSeq` introduces an ORF-aware, quantitative framework for ribosome profiling, delivering end-to-end workflows for ORF annotation, read summarisation, contrast estimation, and visualisation to uncover translational control events at scale.

## 1 INTRODUCTION

Translation is a major regulatory layer of gene expression, yet existing differential translation analyses operate at the gene level and cannot resolve *cis*-regulatory events that occur between open reading frames (ORFs) within the same transcript. Ribosome profiling (Ribo-seq) provides nucleotide-resolution snapshots of ribosome occupancy [1, 2] and has revealed extensive translation of small ORFs, including upstream and downstream ORFs (uORFs and dORFs) [3, 4, 5, 6]. These elements can encode functional micropeptides or modulate main ORF (mORF) translation [7, 5], exemplified by uORF-mediated translational control of ATF4 [8, 9, 10, 11]. However, ORF-level regulation is largely inaccessible to gene-level methods for differential translation [12, 13, 14, 15, 16].

We present `DOTSeq`, a statistical framework for genome-wide ORF-level differential analysis in bulk Ribo-seq with matched RNA-seq, alongside ORF-aware read summarisation for single-cell Ribo-seq (scRibo-seq). `DOTSeq` introduces Differential ORF Usage (`DOU`), which models an ORF’s relative contribution to its gene’s translation output using a beta-binomial generalised linear model (GLM) with sequencing strategy-aware dispersion and an interaction term (`condition:strategy`) that isolates translation-specific effects. To complement `DOU`, `DOTSeq` implements ORF-level Differential Translation Efficiency (`DTE`) using a negative binomial GLM in `DESeq2` [17], capturing monotonic changes in ribosome loading relative to RNA abundance.

Beyond bulk datasets, `DOTSeq` provides ORF-level read summarisation for scRibo-seq, producing ORF-aware count matrices suitable for downstream modelling and integration. This enables users to leverage established single-cell workflows for dimensionality reduction, clustering, and generalised linear mixed modelling (GLMM) using the ORF-level counts generated by `DOTSeq`.

Applied to cell cycle datasets, `DOTSeq` reveals condition-dependent shifts in uORF and mORF usage. Benchmarks show that `DOU` delivers superior sensitivity with robust error control, whereas `DTE` and existing methods are competitive when technical noise is low. Together, these capabilities demonstrate `DOTSeq` as an ORF-aware, quantitative framework for dissecting translational control.

## 2 RESULTS

### 2.1 Overview of DOTSeq modules

Ribo-seq data contain rich information about translation, including complex *cis*-regulatory events such as alternative initiation. These events often manifest as changes in the relative usage of ORFs within the same gene, where standard gene-level differential translation analyses cannot capture. To address this, we developed `DOTSeq`, a statistical framework that enables genome-wide detection of *cis*-translational control. `DOTSeq` uses ORF-level annotations and statistical modelling strategies that capture both absolute changes in translation and condition-specific shifts in the expected proportion of Ribo-seq to RNA-seq reads for individual ORFs, relative to other ORFs within the same gene.

`DOTSeq` assumes preprocessed BAM files generated after adapter trimming and mapping of Ribo-seq and RNA-seq reads with a splice-aware short-read aligner such as `STAR` or `HISAT2` [18, 19] (Figure 1). ORF-level analysis can then proceed via the **Bioconductor ecosystem path**, in which reference GTF and transcript FASTA files are parsed with `DOTSeq::getORFs` to generate discrete, non-overlapping ORF annotations in GRanges format; reads mapping outside exons can be filtered via `getExonicReads`; ORF-level read summarisation for bulk and single-cell data is performed with `countReads` and `countReadsSingleCell`, respectively. Alternatively, an **external flattened-annotation path** flattens reference GTF and transcript FASTA files with the `orf to gtf.py` wrapper (via the `RIBOSS` engine) to produce flattened GTF/BED files containing non-overlapping ORF bins [20]; reads are then summarised with a tool such as `featureCounts` [21]. Both routes flatten ORFs across transcript isoforms into discrete, non-overlapping ORF bins, conceptually similar to `DEXSeq`’s approach to defining exon counting bins [22].

**Figure 1:**
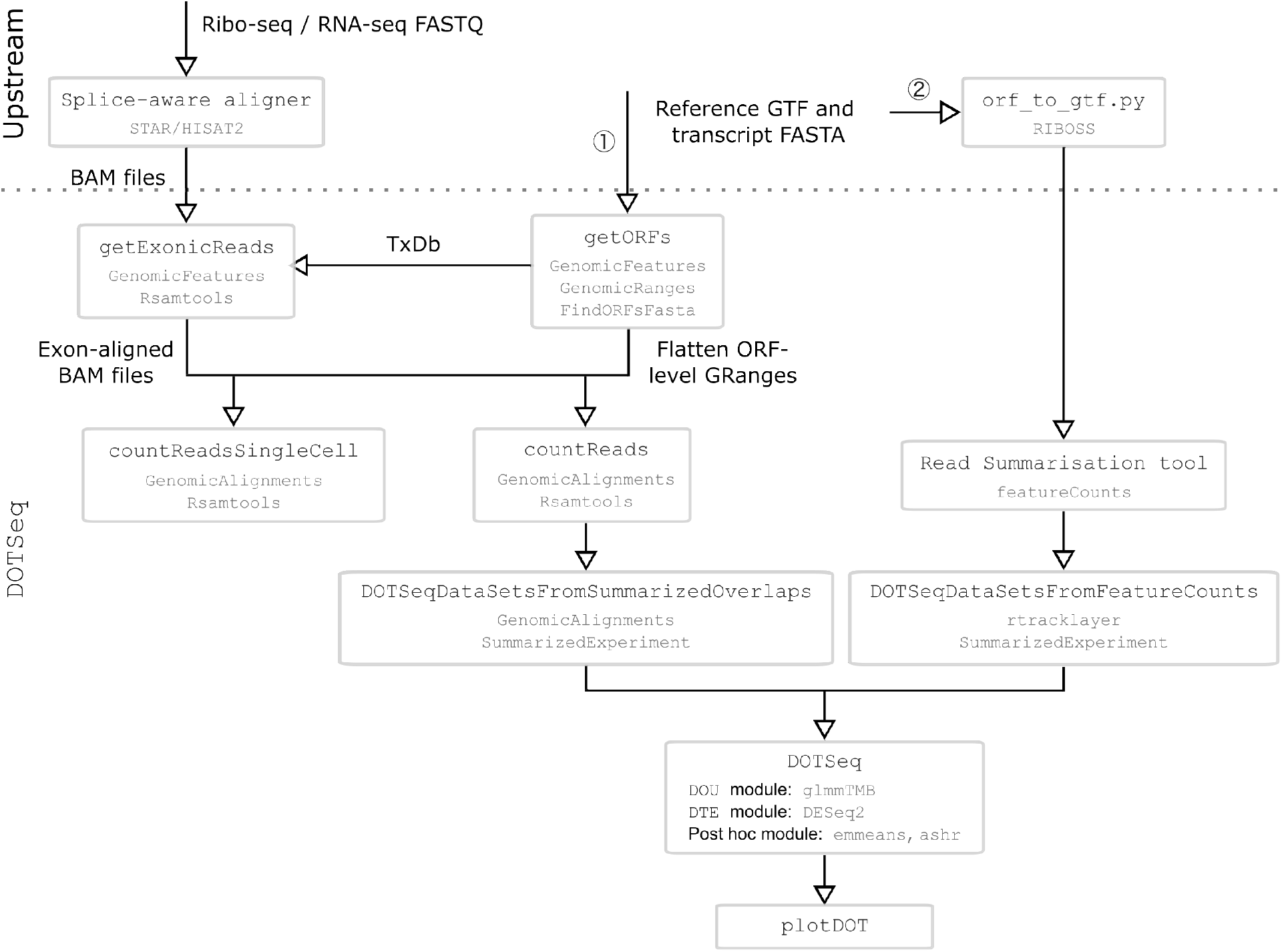
Overview of the DOTSeq workflow. Ribo-seq and/or matched RNA-seq reads are aligned to a reference genome using a splice-aware aligner such as `STAR` or `HISAT2`. ① **Bioconductor ecosystem path:** reference GTF and transcript FASTA files are parsed by `getORFs` to generate discrete, non-overlapping ORF annotations; reads mapping outside exons can be filtered with `getExonicReads`; read summarisation for bulk and single-cell data is performed with `countReads` and `countReadsSingleCell`, respectively, using ORF annotations in GRanges format. ② **External flattened-annotation path:** reference GTF and transcript FASTA files are parsed with the `orf_to_gtf.py` wrapper (via the `RIBOSS` engine) to produce flattened GTF/BED files with ORF-level annotations; reads are then summarised with a tool such as `featureCounts`. Next, DOTSeqDataSets are prepared and subjected to differential analysis. `DOTSeq` provides two statistical modules: Differential ORF Usage (`DOU`) and Differential Translation Efficiency (`DTE`). `plotDOT` generates visualisations for DOU and DTE results.

`DOTSeq` comprises two statistical modules to model differential usage and translation of ORFs (Table 1). The Differential ORF Usage (`DOU`) module captures changes in relative usage of ORFs within a gene to identify *cis*-regulatory events. It uses a beta-binomial GLM implemented via the `glmmTMB` package [23]. Specifically, `DOU` models changes in the expected proportion of Ribo-seq and RNA-seq counts for a given ORF relative to the total counts from other ORFs in the same gene, and tests whether this proportion changes between conditions using an interaction term (`condition:strategy`). This term captures whether the effect of sequencing strategy (Ribo-seq vs RNA-seq) differs between biological conditions, enabling detection of condition-specific shifts in ORF usage driven by *cis*-translation control. For example, if an ORF’s usage remains stable across conditions, its expected proportion of Ribo-seq and RNA-seq counts relative to other ORFs will be consistent, and no condition-specific change in its relative contribution to the gene’s overall expression and translation will be detected. However, if an ORF becomes more or less translated in one condition, its expected Ribo-seq proportion will shift relative to its expected RNA-seq proportion and to other ORFs in the gene, indicating a change in ORF usage within the gene context. For this module, dispersion is modelled separately for Ribo-seq and RNA-seq to account for distinct sources of variability. Post hoc analysis includes pairwise contrasts across conditions, computation of estimated marginal means (EMMs) using the `emmeans` package [24], and adaptive shrinkage of effect sizes via the `ashr` package [24].

**Table 1:**
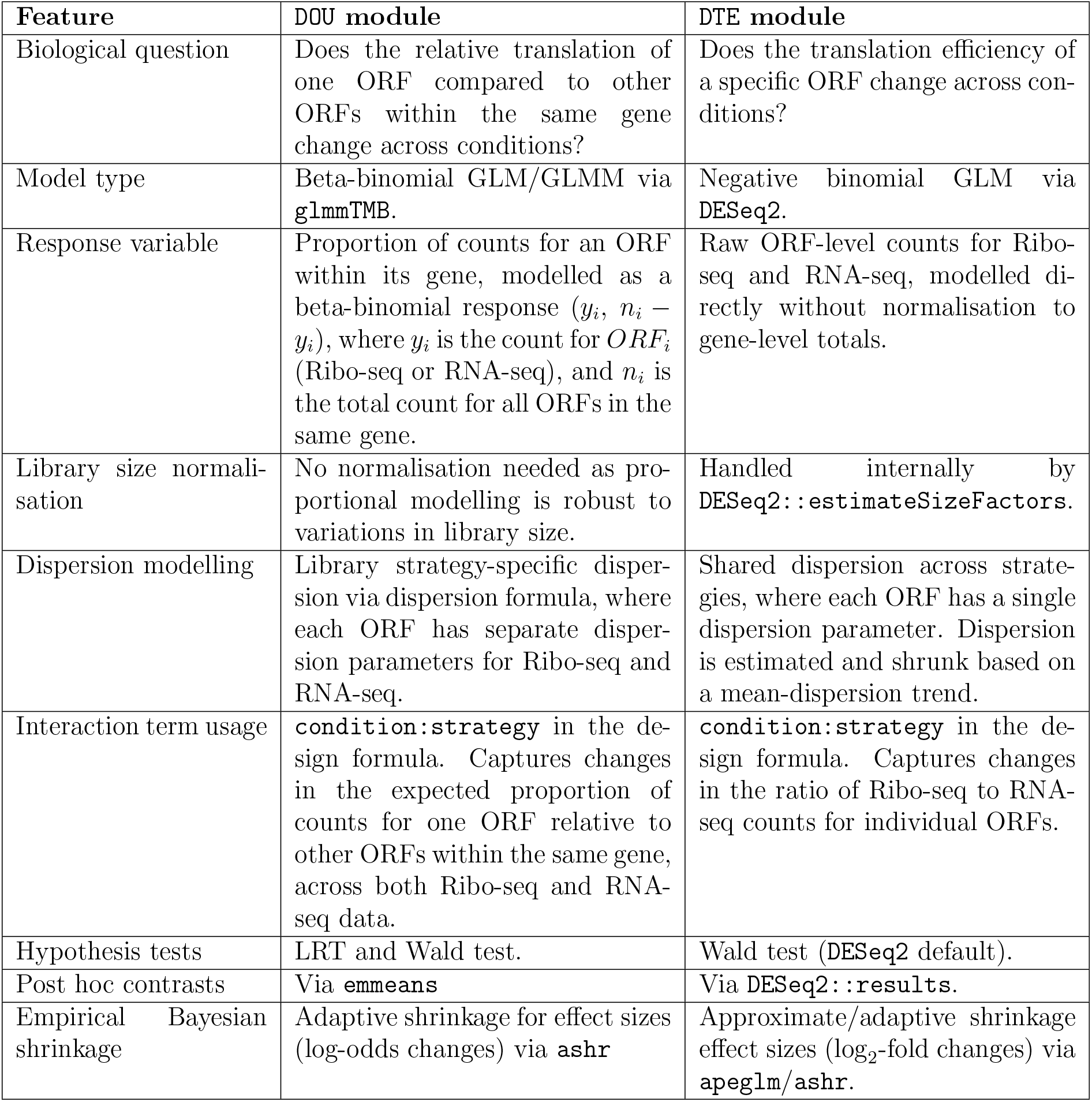
DOTSeq statistical modules for Differential ORF Usage (DOU) and Differential Translation Efficiency (DTE).

In contrast, the Differential Translation Efficiency (`DTE`) module quantifies changes in ribosome loading relative to RNA expression at the level of individual ORFs. It is designed to detect DTE in single-ORF/monocistronic genes and in ORFs showing monotonic changes within multi-cistronic genes that the `DOU` module is not designed to capture. The `DTE` module is conceptually equivalent to `deltaTE` [16] applied at the ORF level. This module models raw ORF-level counts directly using a negative binomial generalised linear model (GLM) implemented in `DESeq2` [17], and tests whether the ratio of Ribo-seq to RNA-seq counts differs across conditions using an interaction term (`condition:strategy`). Together, these complementary modules offer a genome-wide approach for extracting regulatory information in Ribo-seq experiments across biological conditions, enabling systematic interpretation of complex *cis*-regulatory events that are inaccessible to current gene-level tools.

### 2.2 DOTSeq DOU reveals cell cycle-dependent shifts in ORF usage not captured by DTE

To demonstrate the utility of `DOTSeq`, we analysed publicly available bulk Ribo-seq and matched RNA-seq data for HeLa cells synchronised at three distinct cell cycle stages: mitotic cycling (cells undergoing DNA replication and mitosis), mitotic arrest, and interphase (cells replicating DNA without mitosis) [25]. The DOU analysis revealed changes in ORF usage within genes that would be missed by a DTE approach alone. In particular, we observed a significant shift towards increased uORF usage in mitotic cycling versus interphase (Figure 2A). Such differences highlight the impact of *cis*-regulation, as uORF translation often represses mORFs.

**Figure 2:**
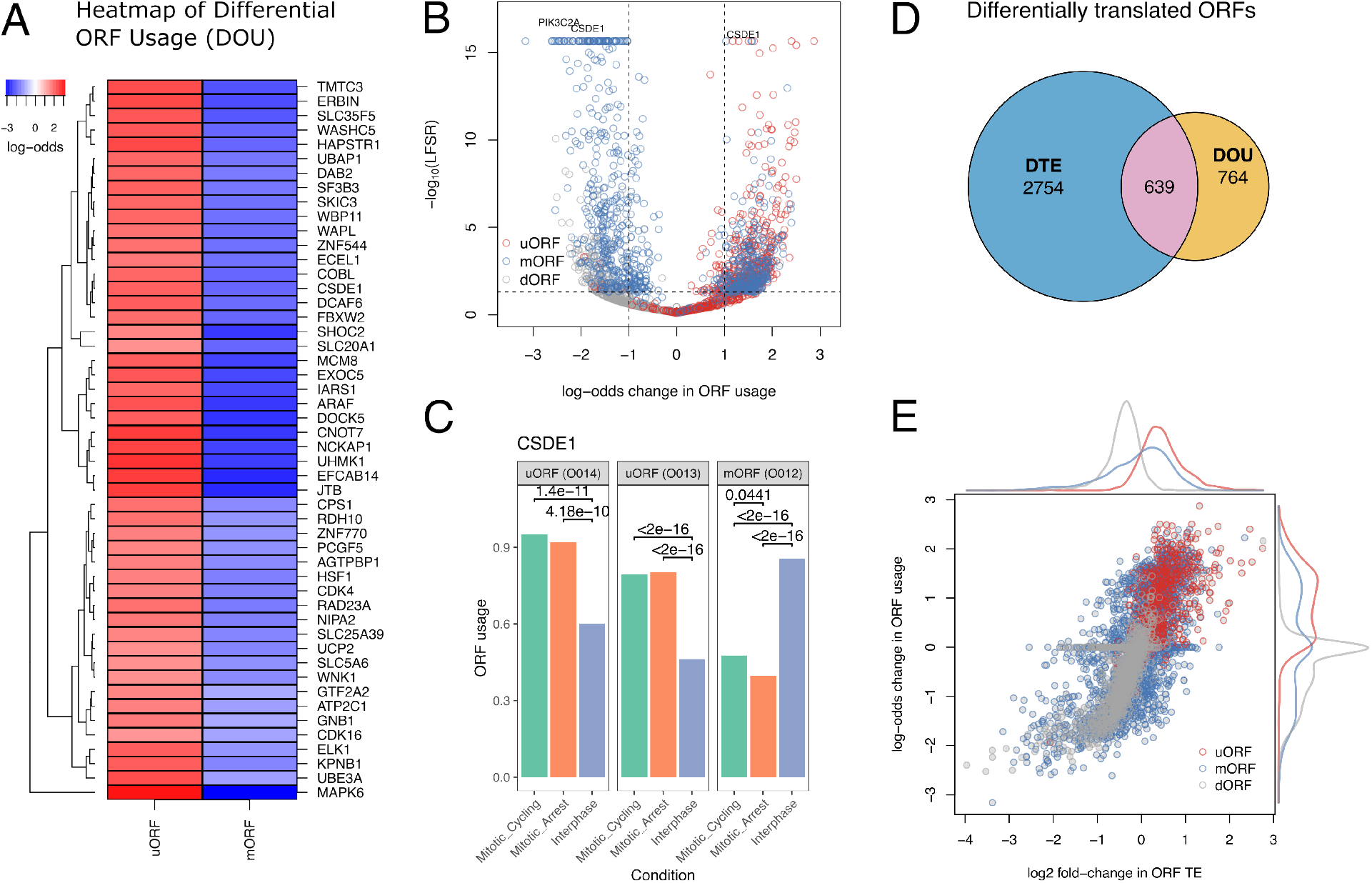
DOTSeq reveals distinct patterns of differential ORF translation between mitotic cycling and interphase. Analysis of Ribo-seq and RNA-seq data comparing HeLa cells synchronised at mitotic cycling, mitotic arrest, and interphase. **(A)** Heatmap of top 50 genes with DOU, showing log-odds values ranging approximately from −3 to +2. Rows represent genes, highlighting differences in uORF and mORF usage between mitotic cycling and interphase conditions. **(B)** Volcano plot of DOU results. The ORFs are plotted by effect sizes (log-odds ratio) and significance [− log_10_ local false sign rate (LFSR)], coloured by ORF type. The top hits are annotated with the gene symbols; the top hit in this analysis is CSDE1. **(C)** The ORF usage plot for the top hit gene, CSDE1, showing the estimated ORF usage probabilities across conditions. **(D)** Venn diagram illustrating overlap between ORFs detected by the `DOU` and `DTE` modules. Of the total, 2754 ORFs are unique to DTE, 764 to DOU, and 639 are shared, yielding an overlap coefficient of 0.46. Plots were generated using `DOTSeq::plotDOT`. **(E)** Scatter plot showing a significant positive correlation between DOU log-odds changes and DTE log-fold changes (Spearman *ρ* = 0.72, *p*-value < 2 × 10^−16^). Effect sizes for both DOU and DTE have been shrunk using empirical Bayesian estimators to improve stability and interpretability.

These significant changes include a marked increase uORF usage and/or decrease in mORF usage in the regulatory-associated protein of mTOR [raptor] (RPTOR), Mitogen-Activated Protein Kinase 6 (MAPK6), Jumping Translocation Breakpoint (JTB), EF-Hand Calcium Binding Domain Containing 14 (EFCAB14), and CCR4-NOT Transcription Complex Subunit 7 (CNOT7) genes (Supplementary Table S1, Figure 2A). This pattern is consistent with previous studies, showing that global translation is repressed during mitosis, at least partly to a greater extent than transcription [26, 27, 28, 29]. While this repression has largely been attributed to phosphorylation-dependent regulation of components of the translational machinery [27, 30, 31, 32], our findings are consistent with an additional layer of translational control mediated by uORFs. In contrast to mitotic cycling, RPTOR and EIF4G1 mORF usage is significantly elevated during interphase (Supplementary Table S1, Figure 2A). These observations align with our understanding that raptor and eIF4G1 are two master regulators of protein synthesis [33, 34]. Raptor is part of the mTORC1 complex that promotes protein synthesis, cell growth and metabolism [33, 35], whereas eIF4G1 is part of the cap-binding complex that mediates recruitment of ribosomes to mRNAs [36]. Together, these patterns are consistent with uORF-mediated control of mORF translation during the cell cycle, particularly for regulators of protein synthesis (e.g., RPTOR, EIF4G1).

A similar trend was observed for MAPK6, also known as ERK3, which is an atypical MAP kinase involved in regulating mitotic progression through signaling pathways controlling cell cycle checkpoints, mitotic entry, and exit. ERK3 is known to become hyper-phosphorylated at multiple sites within its C-terminal domain during mitosis, which increases its protein stability [37]. In our dataset, we observed increased uORF usage in MAPK6 during the mitotic cycling phase, suggesting a potential regulatory mechanism limiting mORF translation. This may help control ERK3 protein levels during mitosis, preventing excessive accumulation that could otherwise accelerate mitotic progression or disrupt cell cycle checkpoints.

To visualise ORFs with strong differential ORF usage, a volcano plot was constructed showing the relationship between the estimated effect sizes (log-odds ratio) and statistical significance (− log_10_ local false sign rate, LFSR). The top hits identified in this analysis were the Cold Shock Domain-Containing Protein E1 (CSDE1) and Phosphatidylinositol-4-phosphate 3-kinase catalytic subunit type 2 alpha (PIK3C2A)

(Figure 2B). To further investigate these signals, gene-level ORF usage plots were generated. These plots show that CSDE1 contains two uORFs that are significantly more utilised than the main ORF (mORF) during both mitotic arrest and mitotic cycling, whereas the mORF is preferentially used during interphase (Figure 2C). This pattern suggests that uORFs may regulate CSDE1 mORF translation across different cell cycle states. Given the role of CSDE1 in tumor progression and immune regulation, and its recent identification as a potential prognostic biomarker and therapeutic target in cancer, these findings illustrate how DOU analysis can reveal shifts in ORF usage within genes across conditions [38, 39, 40, 41, 42, 43]. Such insights highlight the potential of uORF-mediated regulation as an endogenous mechanism to control CSDE1 expression and may inform future translational research.

To assess whether the `DOU` and `DTE` modules capture overlapping or distinct aspects of translational control, we compared effect sizes between the two modules using the HeLa cell cycle data above. Most changes in translation efficiency fell within a modest range (±1 log_2_-fold change), where 764 differentially translated ORFs were missed by the `DTE` module but detected by `DOU` (Figure 2D and 2E, Supplementary Tables S1 and S2; significant threshold of 0.05). However, there was a strong correlation between DOU log-odds changes and DTE log-fold changes, indicating that both modules detect condition-dependent changes in translation through different modelling approaches (Figure 2E, Spearman *ρ* = 0.72, *p*-value < 2 × 10^−16^). Notably, the overlap between the ORFs meeting statistical thresholds was partial: 2754 ORFs were unique to DTE, 764 unique to DOU, and 639 shared (Figure 2D, overlap coefficient = 0.46). These findings demonstrate that the two modules provide complementary insights rather than redundant results.

### 2.3 DOTSeq uncovers cell cycle-dependent translational control at single-cell resolution

Next, we analysed a recently published scRibo-seq dataset for hTERT-RPE-1 cells expressing FUCCI reporters to examine how translation is modulated during cell cycle at single-cell resolution. We performed ribosome footprint summarisation using `DOTSeq::countReadsSingleCell` and modelled the expected per-cell proportion of footprints mapped to uORF [*u/*(*u* + *m*); where u and m indicate total uORF and mORF footprints per cell] with a beta-binomial GLMM, including batch as a random intercept. Model diagnostics indicated zero-deflation expected from filtering, no dispersion issue, and a residual uniformity deviation.

Cells in G0 and mitosis showed a higher per-cell footprints mapping to uORFs relative to mORFs (Figure 3), indicating increased ribosome occupancy at uORFs. The EMMs on the probability scale (Figure 3A) summarise the model-adjusted uORF fraction by treatment, and the Tukey-adjusted, response-scale pairwise contrasts (percentage-point differences; Figure 3B) highlight the magnitude of the shifts between cell cycle stages.

**Figure 3:**
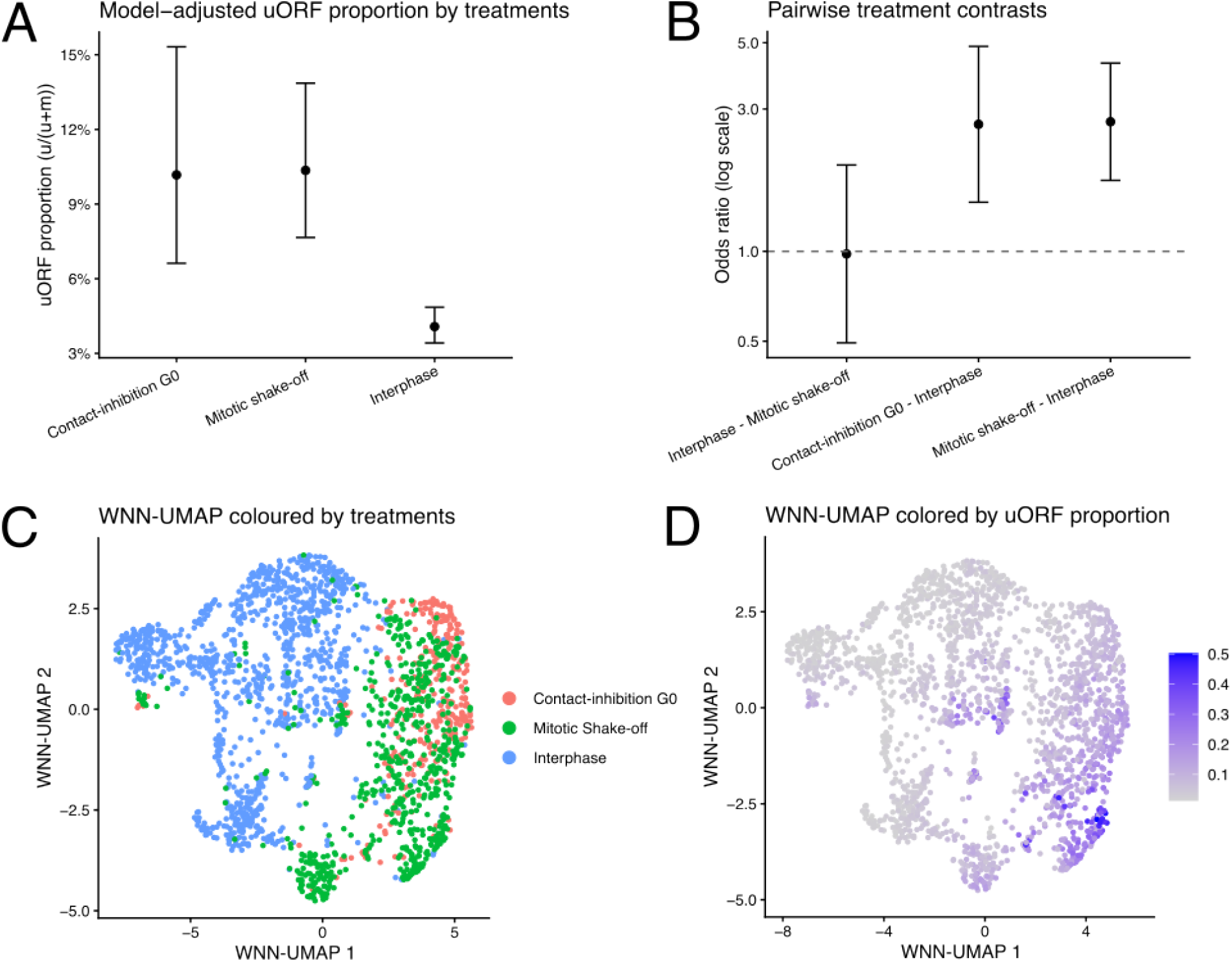
DOTSeq reveals cell cycle-specific translational control in single-cell Ribo-seq (scRibo-seq). **(A)** Model-adjusted uORF fraction by treatments. Points are estimated marginal means (EMMs) on the probability scale; error bars indicate 95% CIs (back-transformed from the logit). **(B)** Pairwise treatment contrasts shown as odds ratios (log scale) with Tukey-adjusted 95% CIs. **(C)** WNN-UMAP (Weighted Nearest Neighbour UMAP) coloured by treatment. **(D)** WNN-UMAP coloured by the per-cell uORF fraction *u/*(*u* + *m*), where *u* and *m* are total uORF and mORF footprints per cell, respectively. All model outputs in **(A** and **B)** are from a beta-binomial generalised linear mixed model (GLMM) for *u/*(*u* + *m*) with a random intercept for batch.

We compared UMAPs derived from the mORF-only, uORF-only, and joint (mORF+uORF) spaces. The joint manifold (Figure 3C and D) showed clearer localised treatment enrichment and pockets of higher uORF usage than either single-modality map (Supplementary Figure S1), consistent with the differential ORF usage in bulk Ribo-seq observed above (Figure 2). Communities retained weak-moderate concordance with treatment (adjusted mutual information, AMI 0.25; permutation *p* ≈ 0.002), indicating that treatment contributes to, but does not define, footprinting-based communities. Together, these patterns support the observations where uORFs are involved in regulating translation across cell cycle stages.

### 2.4 DOTSeq DOU outperforms existing methods across effect sizes

To evaluate performance under controlled conditions, we designed simulation scenarios modelling DOU using `DOTSeq::simDOT`, which builds on the `polyester` simulation engine [44]. These scenarios incorporated experimentally derived Ribo-seq and matched RNA-seq count matrices from the HeLa cell cycle dataset above [25]. To assess sensitivity across biologically relevant effect sizes (Figure 2B), we introduced varying magnitudes of log_2_ changes to multi-cistronic genes by applying opposite-signed effects to uORFs and mORFs. Specifically, we varied *g* ∈ {0.5, 1.0, 1.5, 2.0}, yielding absolute within-gene differences of 2*g* ∈ {1, 2, 3, 4} log_2_-fold (Figure 4, 5). Batch effects were included to provide realistic experimental variability.

**Figure 4:**
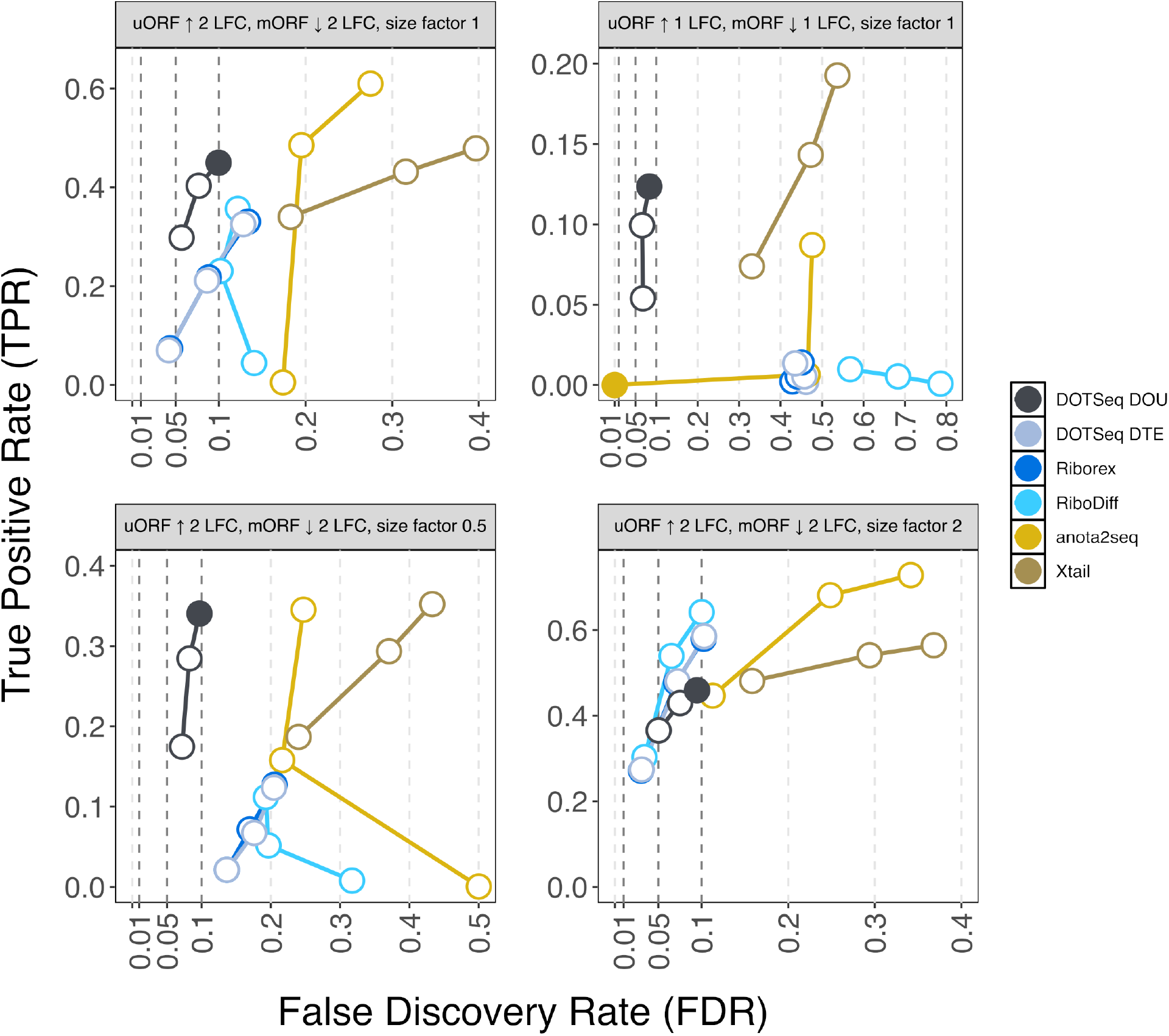
DOTSeq detects simulated *cis*-translational control across effect sizes and size factors. FDR-TPR curves compare `DOTSeq`’s `DOU` and `DTE` modules with existing methods across effect sizes and size factors. Each simulation used Ribo-seq with matched RNA-seq across two conditions, three batches, and one biological replicate per condition (see Methods); batch effects were included to emulate experimental variability. Points show performance at nominal FDRs of 0.01, 0.05, and 0.1, and are filled when the observed FDR is below the nominal threshold. `DOU` points lie closest to the top-left corner, indicating higher TPR at a given observed FDR than alternative methods. Results for `deltaTE` are omitted because they are identical to `DTE` in our benchmarks.

**Figure 5:**
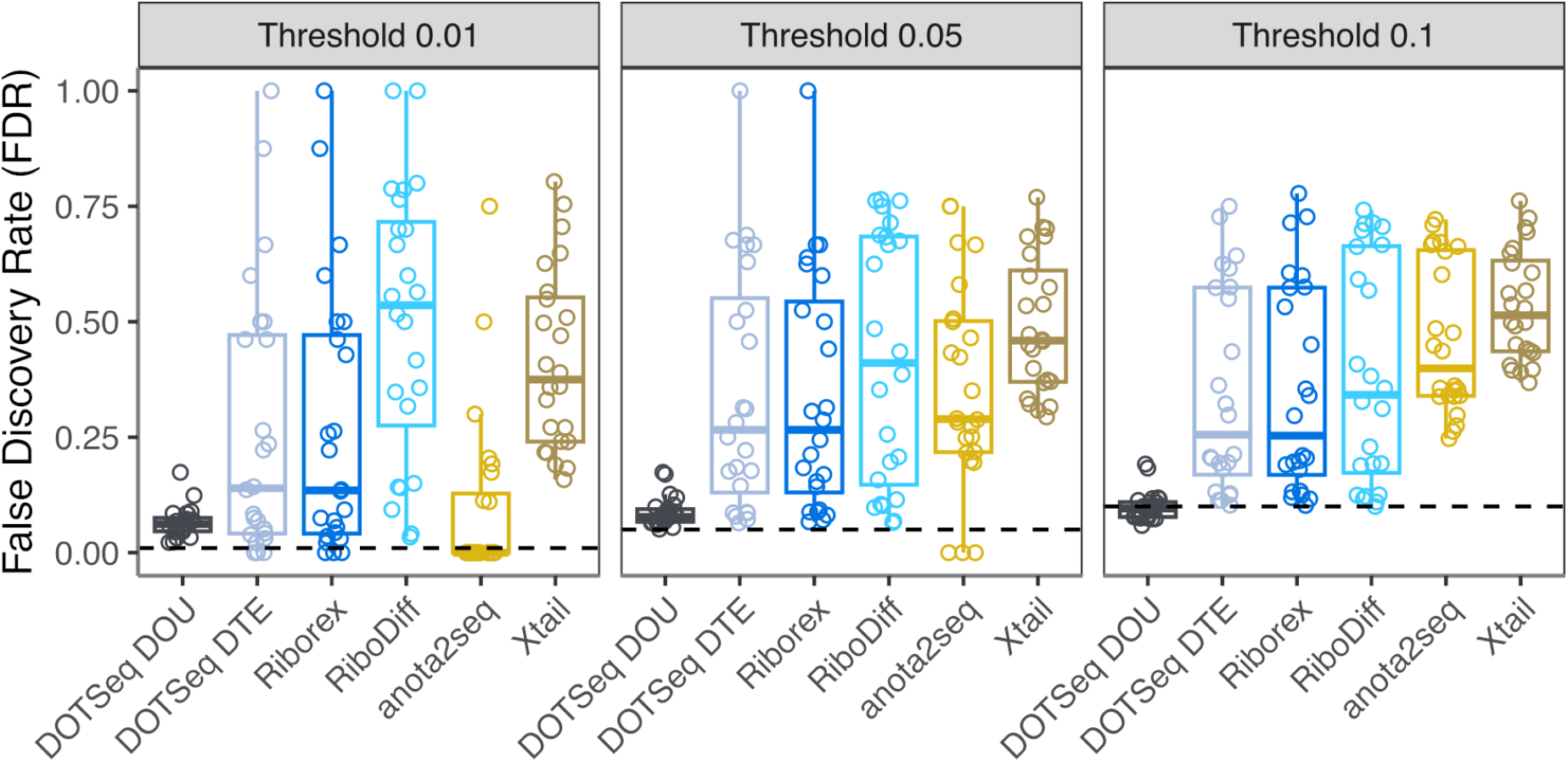
DOTSeq DOU module shows the most consistent and conservative control of FDR for differential ORF translation analysis. The performance of the `DOTSeq DOU` and `DTE` modules was benchmarked against existing tools at nominal FDR thresholds of 0.01, 0.05, and 0.1. Each point in the box plots represents the observed FDR at a given nominal threshold, while the y-axis shows the actual FDR.

We analysed each simulation using the `DOU` and `DTE` modules and benchmarked them against existing methods for differential translation analysis, including `anota2seq` [12], `deltaTE` [16], `RiboDiff` [14], `Riborex` [15], and `Xtail` [13]. Performance was assessed with `iCOBRA`, summarising False Discovery Rate (FDR) and True Positive Rate (TPR) across multiple thresholds [45].

`DOU` outperformed `DTE` and other methods across the tested effect sizes; only at `size_factor`= 2 did some competitors surpass it in sensitivity (Figure 4). Among methods tested, `DOU` was the only one to meet the nominal FDR of 0.1 across settings. These findings emphasise the value of modelling condition-specific proportional changes for an ORF relative to other ORFs within the same gene via a beta-binomial framework. In addition, they highlight the complementary utility of the two `DOTSeq` modules across technical noise levels.

To further assess statistical calibration, we examined the distribution of significance measures produced by the `DOTSeq` modules across the effect-size grid and both regulatory patterns (uORF up/mORF down and uORF down/mORF up). The `DOU` module produced a right-skewed p-value distribution, consistent with well-calibrated significance (Figure 6). In contrast, when applied to DOU-type regulatory events, the `DTE` module exhibited a U-shaped distribution, suggesting p-value inflation and reduced calibration. These results underscore the importance of DOU-specific modelling for detecting differential ORF usage.

**Figure 6:**
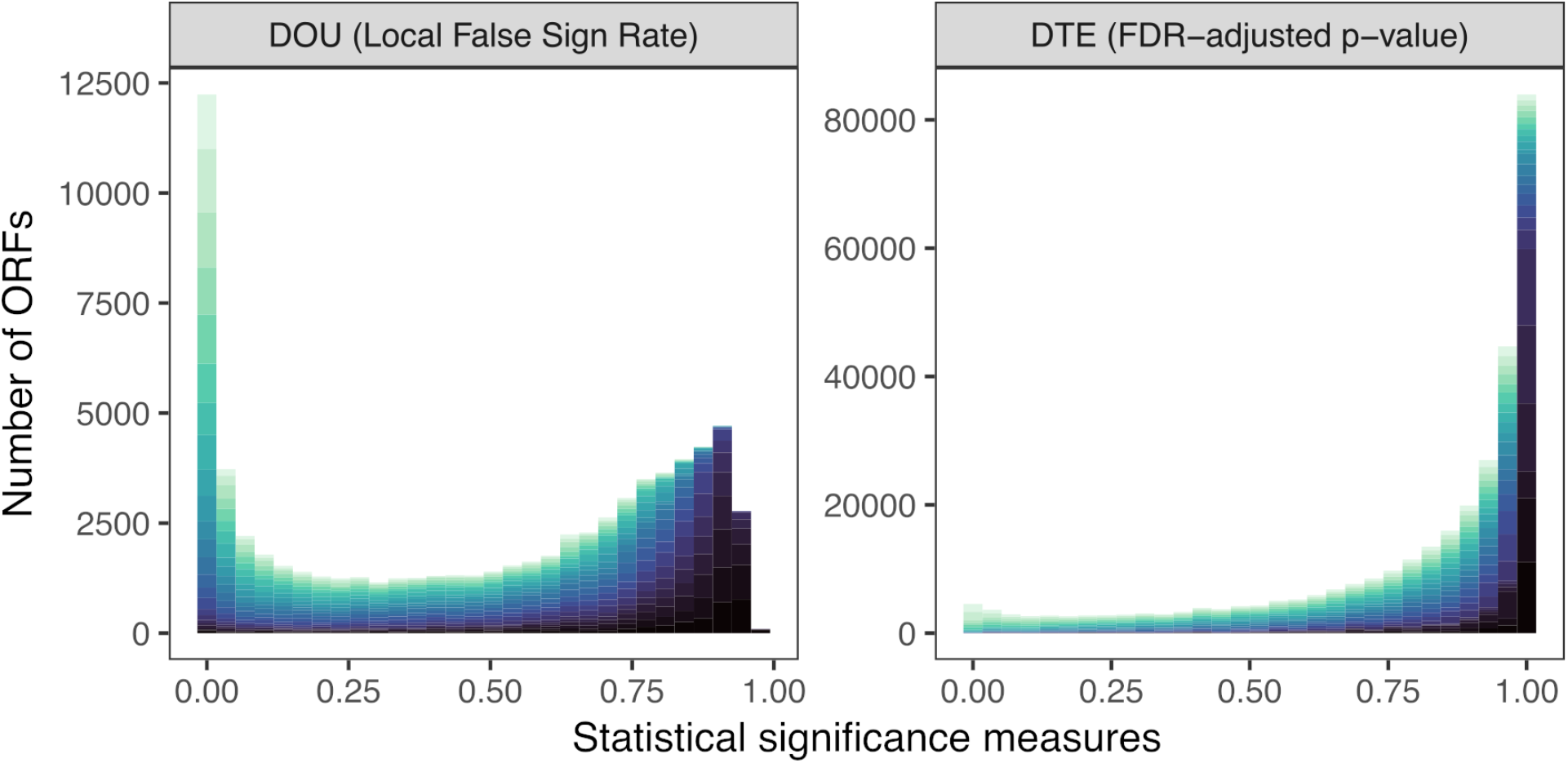
DOTSeq DOU module is well-calibrated for differential ORF usage analysis. Distributions of aggregated Local False Sign Rate (LFSR) and FDR-adjusted p-values across 128 simulated scenarios for the `DOU` and `DTE` modules, respectively. Bar height indicates the total number of ORFs across all scenarios, whereas colour gradient indicates the composition of scenarios contributing to the corresponding p-value bin.The `DOU` module shows a right-skewed distribution, consistent with well-calibrated statistical significance. In contrast, the `DTE` module exhibits a U-shaped distribution, indicating p-value inflation.

## 3 DISCUSSION

The ability to resolve changes in ORF usage across biological conditions opens new avenues for dissecting translational control mechanisms that are obscured by gene-level analyses. `DOTSeq` addresses this need by enabling differential analyses of ORF usage and translation efficiency. Simulations demonstrate `DOU` as a leading approach for ORF-level translational control, combining high sensitivity with near-nominal FDR across effect sizes. This sensitivity and resolution enable the discovery of regulatory events (e.g., uORF-mediated repression) that are often overlooked in gene-level analyses.

Empirically, we identified *cis*-regulatory patterns consistent with widespread translational repression during mitosis, including signatures of uORF-mediated repression of EIF4G1, a master regulator of protein synthesis. Using `DOTSeq`’s single-cell ORF-level summarisation, we also detected a significant increase in per-cell uORF occupancy in mitosis relative to interphase, demonstrating applicability to single-cell datasets and enabling downstream modelling at single-cell resolution.

These discoveries rely on `DOTSeq`’s ability to statistically model overdispersed and sparse count data from Ribo-seq and RNA-seq using the state-of-the-art TMB (Template Model Builder) architecture [23]. However, fitting beta-binomial GLMs on sparse Ribo-seq data can lead to convergence issues, including a non-positive definite Hessian. To mitigate these issues, `DOTSeq` supports a brute force optimisation step using three optimisers implemented in `glmmTMB`: `nlminb`, `bobyqa`, and `optim`. In addition, we have implemented multiple dispersion modelling approaches within our framework. (i) The default approach models dispersion as a function of sequencing strategy (Ribo-seq with RNA-seq as reference), allowing for strategy-dependent overdispersion. Although this approach performs well in simulations, it can be more prone to convergence issues than simpler dispersion models. (ii) As an alternative, users may assume dispersion to be constant across all predictor levels. This approach reduces model complexity and is less prone to convergence issues. (iii) To balance model stability and flexibility, both models are fitted and compared using likelihood ratio tests. The simpler model is selected when no significant difference is detected. (iv) For maximum flexibility, users may specify a custom formula for dispersion modelling. Random effects can be included in the design formula and dispersion formulas within the beta-binomial GLMM framework. Users may also enable model diagnostics to assess model fit, including tests for overdispersion, zero inflation, and residual properties.

While flexible, fitting fully parameterised models comes with a computational cost. For example, it took 6 min 53 s (± 42 s) to fit the beta-binomial GLMs for 35,863 ORFs in the HeLa cell cycle dataset, running on a single thread of a MacBook Pro M4 with 16 GB RAM. In contrast, the `DTE` module runs at a speed comparable to existing tools such as `deltaTE` (31 ± 2 s). To mitigate this computational cost, `DOTSeq` supports parallel processing via `OpenMP`, which is used internally in `glmmTMB` to accelerate model fitting. Altogether, `DOTSeq` offers a flexible and powerful framework for dissecting the complex landscape of translational control. `DOTSeq` is broadly applicable to studies of translational regulation across conditions within biological contexts such as development, disease, and stress responses.

## 4 METHODS

### 4.1 Data

The Ribo-seq and matched RNA-seq data for HeLa cells synchronised at interphase, mitotic arrest, and mitotic cycling were retrieved from PRJNA957808, with two biological replicates for each condition [25]. The single-cell Ribo-seq dataset was obtained from PRJNA680481, consisting of contact-inhibition G0, mitotic shake-off, and interphase sorted fractions for hTERT RPE-1 cells[46].

### 4.2 Data preprocessing

Adapter trimming was first performed on both bulk and single-cell datasets using `Cutadapt` v5.1 [47]. Reads were mapped to the UCSC Genome Browser’s hg38 with GENCODE v47 annotation using `STAR` v2.7.11b [18, 48, 49].

To generate discrete, non-overlapping ORF annotations for read summarisation, high-confidence reference GTF and transcript FASTA files (MANE GRCh38 v1.4) were parsed with `DOTSeq::getORFs`.

For the bulk dataset, the ORF-level quantification was performed using `DOTSeq::countReads`. For the single-cell Ribo-seq dataset, the ORF-level quantification was performed using `DOTSeq::countReadsSingleCell`, which also aggregates counts into sparse matrices and stores the mORF and uORF assays in a `Seurat` v5.5 object for downstream analyses [50].

### 4.3 Statistical modelling for scRibo-seq

The per-cell uORF proportion was calculated as *u/*(*u* + *m*), where *u* and *m* are the total uORF and mORF footprints per cell, respectively, and used as a feature for downstream analyses. For all three UMAPs derived from the mORF-only, uORF-only, and joint (mORF+uORF) spaces, the mORF and uORF counts were normalised, scaled, and subjected to principal component analysis (PCA), nearest-Neighbour graph construction, clustering, and uniform manifold approximation and projection (UMAP) dimensionality reduction. For uORF counts, only cells with more than 50 uORF UMIs were included to reduce noise, although users can adjust this threshold. For the joint manifold analysis, the weighted nearest-Neighbour (WNN) strategy was implemented by integrating mORF and uORF principal components to produce a joint UMAP embedding.

Statistical modelling of the per-cell uORF fraction was performed using a beta-binomial GLMM implemented in `glmmTMB` v1.1.12 [23], with a random intercept for batch (SRR accessions). Estimated marginal means (EMMs) were computed using `emmeans` v1.11.2-8, and pairwise treatment contrasts were expressed as odds ratios. Model diagnostics were performed using `DHARMa` v0.4.7.

### 4.4 Statistical modelling for DOU

The `DOTSeq` framework builds on a beta-binomial GLM implemented in `glmmTMB` v1.1.12 [23]. The expected proportion of reads mapped to an ORF within its gene is modeled by a mean model and a dispersion model. The mean model uses the logit link function to model the expected probability (*µ*_*i*_) that a read maps to a particular ORF for observation *i*.

#### 4.4.1 Testing the interaction effect

`DOTSeq` provides an optional test to assess the significance of a translation-specific effect. A likelihood ratio test can be performed by comparing a full model, which includes an interaction between condition and strategy, against a null model, which contains only additive effects:

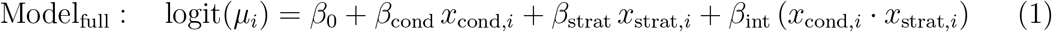

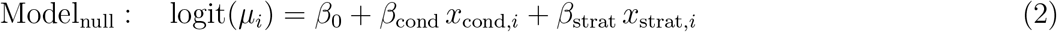

The test compares the log-likelihoods of the full and null models to assess whether the inclusion of the interaction term, *β*_int_, significantly improves model fit.

The unified mean model (Equation 1) can be simplified by substituting the dummy variable values to isolate the effective intercept and slope for each sequencing level. For the RNA-seq level (*x*_strat,*i*_ = 0), the model simplifies to:

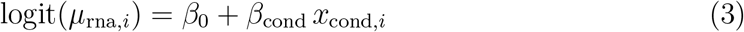

For the Ribo-seq level (*x*_strat,*i*_ = 1), the model simplifies to:

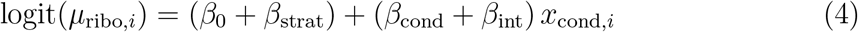

The term *β*_int_ captures the differential effect of condition between the Ribo-seq and RNA-seq levels. Therefore, when *β*_int_ ≠ 0, the condition affects Ribo-seq differently than RNA-seq, indicating a translation-specific effect (DOU).

#### 4.4.2 Contrasts

Since `DOTSeq` models the expected proportion of reads, a condition-dependent contrast between Ribo-seq and RNA-seq levels is performed using `emmeans` v1.11.2-8 [51] with Wald test to isolate these translation-specific effects:

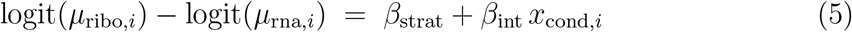

This contrast quantifies how much more (or less) likely a read is to map to an ORF at the Ribo-seq level compared to the RNA-seq level, condition-dependently. Adaptive empirical Bayesian shrinkage is applied to the log-odds contrasts using `ashr` v2.2-63 [24] to stabilise effect size estimates. While not equivalent to DTE, DOU reflects a condition-dependent changes in ORF usage across sequencing strategies, capturing *cis*-regulatory effects on translation.

#### 4.4.3 Dispersion modelling

To account for overdispersion in proportional modelling, `DOTSeq` models variability using the dispersion model in a log link function:

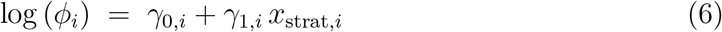

where *ϕ*_*i*_ is the overdispersion parameter for *ORF*_*i*_, *γ*_0_ is the dispersion intercept, and *γ*_1_ captures the additional dispersion at the Ribo-seq level. This allows the model to fit a shared mean structure across Ribo-seq and RNA-seq levels and estimate separate dispersion parameters for these two levels. The fitted dispersions for *ORF*_*i*_ are computed as:

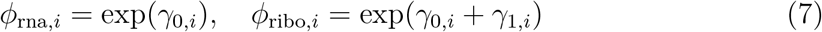

where *γ*_0,*i*_ and *γ*_1,*i*_ are the intercept and strategy effect for *ORF*_*i*_, respectively. This approach accommodates varying technical noise and biological heterogeneity across the sequencing levels, thereby improving statistical inference accuracy.

### 4.5 Statistical modelling for DTE

The `DTE` module in `DOTSeq` uses the `DESeq2` framework to model DTE across conditions directly through the `condition:strategy` interaction term [17, 16]. For each ORF, raw Ribo-seq and RNA-seq read counts are normalised and modelled using a negative binomial GLM. This includes a standard `DESeq2` workflow: size factor estimation, dispersion fitting, Wald tests for condition-specific contrasts, and shrinkage of effect sizes. This complements the `DOU` module by identifying ORFs with overall changes in translation, using transcript abundance as baseline.

### 4.6 Simulation

To simulate *cis*-translational control, 128 datasets were generated using the `DOTSeq` sim-DOT function, which is based on the `deltaTE` source code [16] and uses the `polyester` v1.29.1 engine [44]. The HeLa cell cycle count matrices were used as input to preserve realistic baseline expression and dispersion profiles.

We crossed two predefined regulatory patterns, “uORF up mORF down” and “uORF down mORF up”, with a grid of log_2_ effect sizes *g* ∈ {0.5, 1.0, 1.5, 2.0}. Within each affected gene, uORF and mORF effects were applied with opposite signs, yielding absolute within-gene differences of 2*g* ∈ {1, 2, 3, 4} log_2_-fold. In the simulator, these were passed as natural-log effects by multiplying *g* by log(2). We additionally varied two parameters that modulate dispersion: `size_factor` ∈ {0.5, 1, 2, 4}, which multiplicatively scales the negative-binomial size parameter (*r*; larger values reduce dispersion/technical noise), and `min_size` ∈ {0.5, 1, 2, 4}, which sets a lower bound on *r* to cap extreme dispersion. This grid yielded 2 × 4 × 4 × 4 = 128 scenarios.

Each simulation contained two experimental conditions, one biological replicate per condition (`num_samples` = 1), and three batches (num batches = 3). Batch effects were introduced according to simDOT default settings (applied to a subset of ORFs; mean shift ≈ 0.9 on the log scale) to emulate realistic between-batch variability. For each scenario we produced combined Ribo-seq and RNA-seq count matrices, sample-level metadata (condition, strategy, replicate, batch), ground-truth labels for differentially regulated ORFs, and the corresponding effect sizes.

### 4.7 Benchmarking and analysis

The performance of `DOTSeq` was benchmarked using simulation datasets against four existing tools for detecting differential translation: anota2seq v1.32.0 [12], deltaTE [16], `RiboDiff` v0.2.2 [14], `Riborex` v2.4.0 [15], and `Xtail` v.1.20 [13]. The `DOU` and `DTE` modules of `DOTSeq` were evaluated independently.

Performance was assessed using `iCOBRA` v1.37.1 [45], which was used to generate FDR-TPR curves. The `onlyshared` argument in the `calculate performance` function was set to FALSE as default, allowing all features in the truth table to be used, irrespective of whether they were detected by all methods.

Overlap between significant ORFs identified by the `DOU` and `DTE` modules was visualised using `eulerr` v7.0.2 [52]. Additional plots were produced using the base R functions [53] and `ggplot2` v3.5.2 [45].

### 4.8 Use of LLMs

LLMs were used to generate docstrings, improve clarity, and/or identify inconsistencies in the code and the main text.

### 4.9 Code availability

`DOTSeq` is available at https://bioconductor.org/packages/DOTSeq and https://github.com/compgenom/DOTSeq. Scripts for reproducing the analyses are available at https://github.com/compgenom/DOTSeq_paper_2026.

## 5 AUTHOR CONTRIBUTIONS

C.S.L. conceptualised and led the study, developed the statistical framework and conducted data analysis. G.S.W.C. and C.S.L performed benchmarking analyses, prepared the GitHub repository, and developed the package vignette. C.S.L. and G.S.W.C. wrote the manuscript.

## 6 ACKNOWLEDGEMENTS

The authors appreciate Assoc Prof Chris M. Brown’s feedback on the manuscript.

## References

[1] Nicholas T Ingolia et al. “Genome-wide analysis in vivo of translation with nucleotide resolution using ribosome profiling”. In: science 324.5924 (Apr. 2009), pp. 218–223.

[2] Kotaro Tomuro and Shintaro Iwasaki. “Advances in ribosome profiling technologies”. In: Biochemical Society Transactions (May 2025), BST20253061.

[3] Jonathan M Mudge et al. “Standardized annotation of translated open reading frames”. en. In: Nat Biotechnol 40.7 (July 2022), pp. 994–999.

[4] Eric W Deutsch et al. “High-quality peptide evidence for annotating non-canonical open reading frames as human proteins”. en. Sept. 2024.

[5] Roger P Hellens et al. “The Emerging World of Small ORFs”. en. In: Trends Plant Sci 21.4 (Apr. 2016), pp. 317–328.

[6] Qiushuang Wu et al. “Translation of small downstream ORFs enhances translation of canonical main open reading frames”. en. In: EMBO J 39.17 (Sept. 2020), e104763.

[7] Casimiro Baena-Angulo, Ana Isabel Platero, and Juan Pablo Couso. “Cis to trans: small ORF functions emerging through evolution”. en. In: Trends Genet 41.2 (Feb. 2025), pp. 119–131.

[8] Krishna M Vattem and Ronald C Wek. “Reinitiation involving upstream ORFs regulates ATF4 mRNA translation in mammalian cells”. en. In: Proc Natl Acad Sci U S A 101.31 (Aug. 2004), pp. 11269–11274.

[9] Phoebe D Lu, Heather P Harding, and David Ron. “Translation reinitiation at alternative open reading frames regulates gene expression in an integrated stress response”. en. In: J Cell Biol 167.1 (Oct. 2004), pp. 27–33.

[10] Graham Neill and Glenn R Masson. “A stay of execution: ATF4 regulation and potential outcomes for the integrated stress response”. en. In: Front Mol Neurosci 16 (Feb. 2023), p. 1112253.

[11] Anna M Smirnova et al. “Stem-loop-induced ribosome queuing in the uORF2/ATF4 overlap fine-tunes stress-induced human ATF4 translational control”. en. In: Cell Rep 43.4 (Apr. 2024), p. 113976.

[12] Christian Oertlin et al. “Generally applicable transcriptome-wide analysis of translation using anota2seq”. In: Nucleic acids research 47.12 (2019), e70–e70.

[13] Zhengtao Xiao et al. “Genome-wide assessment of differential translations with ribosome profiling data”. en. In: Nat Commun 7 (Apr. 2016), p. 11194.

[14] Yi Zhong et al. “RiboDiff: detecting changes of mRNA translation efficiency from ribosome footprints”. en. In: Bioinformatics 33.1 (Jan. 2017), pp. 139–141.

[15] Wenzheng Li et al. “Riborex: fast and flexible identification of differential translation from Ribo-seq data”. en. In: Bioinformatics 33.11 (June 2017), pp. 1735–1737.

[16] Sonia Chothani et al. “deltaTE: Detection of Translationally Regulated Genes by Integrative Analysis of Ribo-seq and RNA-seq Data”. en. In: Curr Protoc Mol Biol 129.1 (Dec. 2019), e108.

[17] Michael I Love, Wolfgang Huber, and Simon Anders. “Moderated estimation of fold change and dispersion for RNA-seq data with DESeq2”. en. In: Genome Biol 15.12 (2014), p. 550.

[18] Alexander Dobin et al. “STAR: ultrafast universal RNA-seq aligner”. en. In: Bioinformatics 29.1 (Jan. 2013), pp. 15–21.

[19] Daehwan Kim et al. “Graph-based genome alignment and genotyping with HISAT2 and HISAT-genotype”. en. In: Nat Biotechnol 37.8 (Aug. 2019), pp. 907–915.

[20] Chun Shen Lim et al. “RIBOSS detects novel translational events by combining long- and short-read transcriptome and translatome profiling”. en. In: Brief Bioinform 26.2 (Mar. 2025).

[21] Yang Liao, Gordon K Smyth, and Wei Shi. “featureCounts: an efficient general purpose program for assigning sequence reads to genomic features”. en. In: Bioinformatics 30.7 (Apr. 2014), pp. 923–930.

[22] Simon Anders, Alejandro Reyes, and Wolfgang Huber. “Detecting differential usage of exons from RNA-seq data”. en. In: Genome Res 22.10 (Oct. 2012), pp. 2008–2017.

[23] Maeve McGillycuddy et al. “Parsimoniously fitting large multivariate random effects in glmmTMB”. en. In: J. Stat. Softw. 112.1 (2025).

[24] Matthew Stephens. “False discovery rates: a new deal”. en. In: Biostatistics 18.2 (Apr. 2017), pp. 275–294.

[25] Jimmy Ly et al. “Nuclear release of eIF1 restricts start-codon selection during mitosis”. en. In: Nature 635.8038 (Nov. 2024), pp. 490–498.

[26] Marvin E Tanenbaum et al. “Regulation of mRNA translation during mitosis”. en. In: Elife 4 (Aug. 2015).

[27] Mikhail I Dobrikov et al. “Mitotic phosphorylation of eukaryotic initiation factor 4G1 (eIF4G1) at Ser1232 by Cdk1:cyclin B inhibits eIF4A helicase complex binding with RNA”. en. In: Mol Cell Biol 34.3 (Feb. 2014), pp. 439–451.

[28] A Contreras and C Perea-Resa. “Transcriptional repression across mitosis: mechanisms and functions”. en. In: Biochem Soc Trans 52.1 (Feb. 2024), pp. 455–464.

[29] Qinyu Hao et al. “The S-phase-induced lncRNA SUNO1 promotes cell proliferation by controlling YAP1/Hippo signaling pathway”. en. In: eLife (Oct. 2020).

[30] Dana M Gwinn et al. “AMPK phosphorylation of raptor mediates a metabolic checkpoint”. In: Molecular cell 30.2 (2008), pp. 214–226.

[31] Mohamed Moustafa-Kamal et al. “The mTORC1/S6K/PDCD4/eIF4A axis determines outcome of mitotic arrest”. In: Cell Reports 33.1 (2020).

[32] Richard I Odle et al. “An mTORC1-to-CDK1 switch maintains autophagy suppression during mitosis”. In: Molecular cell 77.2 (2020), pp. 228–240.

[33] Long He, Sungyun Cho, and John Blenis. “mTORC1, the maestro of cell metabolism and growth”. In: Genes & Development 39.1-2 (2025), pp. 109–131.

[34] Nathan R James and John S O’Neill. “Circadian Control of Protein Synthesis”. In: BioEssays 47.3 (2025), e202300158.

[35] Jay N Joshi et al. “mTORC1 activity oscillates throughout the cell cycle, promoting mitotic entry and differentially influencing autophagy induction”. In: Cell Reports 43.8 (2024).

[36] Jailson Brito Querido, Irene Díaz-López, and V Ramakrishnan. “The molecular basis of translation initiation and its regulation in eukaryotes”. In: Nature Reviews Molecular Cell Biology 25.3 (2024), pp. 168–186.

[37] Pierre-Luc Tanguay, Geneviève Rodier, and Sylvain Meloche. “C-terminal domain phosphorylation of ERK3 controlled by Cdk1 and Cdc14 regulates its stability in mitosis”. In: Biochemical Journal 428.1 (Apr. 2010), pp. 103–111. doi: 10.1042/bj20091604.

[38] Ao-Xiang Guo et al. “The role of CSDE1 in translational reprogramming and human diseases”. In: Cell Communication and Signaling 18.1 (Jan. 2020). doi: 10.1186/s12964-019-0496-2.

[39] Jiandong Liu et al. “N6-methyladenosine-modified lncRNA ARHGAP5-AS1 stabilises CSDE1 and coordinates oncogenic RNA regulons in hepatocellular carcinoma”. In: Clinical and Translational Medicine 12.11 (Nov. 2022). doi: 10.1002/ctm2.1107.

[40] Laurence Wurth et al. “UNR/CSDE1 Drives a Post-transcriptional Program to Promote Melanoma Invasion and Metastasis”. In: Cancer Cell 36.3 (Sept. 2019), p. 337. doi: 10.1016/j.ccell.2019.08.013.

[41] Javier Martinez-Useros et al. “UNR/CSDE1 Expression Is Critical to Maintain Invasive Phenotype of Colorectal Cancer through Regulation of c-MYC and Epithelial-to-Mesenchymal Transition”. In: Journal of Clinical Medicine 8.4 (Apr. 2019), pp. 560–560. doi: 10.3390/jcm8040560.

[42] Tian Ningyu et al. “RNA-binding Protein UNR Promotes Glioma Cell Migration and Regulates the Expression of Ribosomal Protein L9”. In: Chinese Medical Sciences Journal 33.3 (2018), pp. 143–151. doi: 10.24920/11815.

[43] Wanting Li et al. “CSDE1 depletion inhibits tumor progression through enhancing B-cell infiltration in NSCLC”. In: Cell Death & Disease 17.1 (Dec. 2025). doi: 10.1038/s41419-025-08282-9.

[44] Alyssa C Frazee et al. “Polyester: simulating RNA-seq datasets with differential transcript expression”. en. In: Bioinformatics 31.17 (Sept. 2015), pp. 2778–2784.

[45] Charlotte Soneson and Mark D Robinson. “iCOBRA: open, reproducible, standardized and live method benchmarking”. en. In: Nat Methods 13.4 (Apr. 2016), p. 283.

[46] Michael VanInsberghe et al. “Single-cell Ribo-seq reveals cell cycle-dependent translational pausing”. In: Nature 597.7877 (Sept. 2021), pp. 561–565. doi: 10.1038/s41586-021-03887-4. url: https://www.nature.com/articles/s41586-021-03887-4.

[47] Marcel Martin. “Cutadapt removes adapter sequences from high-throughput sequencing reads”. In: EMBnet J. 17.1 (May 2011), p. 10.

[48] Gerardo Perez et al. “The UCSC Genome Browser database: 2025 update”. en. In: Nucleic Acids Res 53.D1 (Jan. 2025), pp. D1243–D1249.

[49] Jonathan M Mudge et al. “GENCODE 2025: reference gene annotation for human and mouse”. en. In: Nucleic Acids Res 53.D1 (Jan. 2025), pp. D966–D975.

[50] Yuhan Hao et al. “Dictionary learning for integrative, multimodal and scalable single-cell analysis”. In: Nature Biotechnology (2023). doi: 10.1038/s41587-023-01767-y. url: https://doi.org/10.1038/s41587-023-01767-y.

[51] Russell V. Lenth. emmeans: Estimated Marginal Means, aka Least-Squares Means. R package version 1.11.2-80003. 2025. url: https://rvlenth.github.io/emmeans/.

[52] Johan Larsson. “Area-Proportional Euler and Venn Diagrams with Ellipses [R package eulerr version 7.0.2]”. In: (Mar. 2024).

[53] The R Project for Statistical Computing. en. https://www.R-project.org/. Accessed: 2025-9-21.

