## Supplementary Figure S1 for "DOTSeq enables genome-wide detection of differential ORF usage"

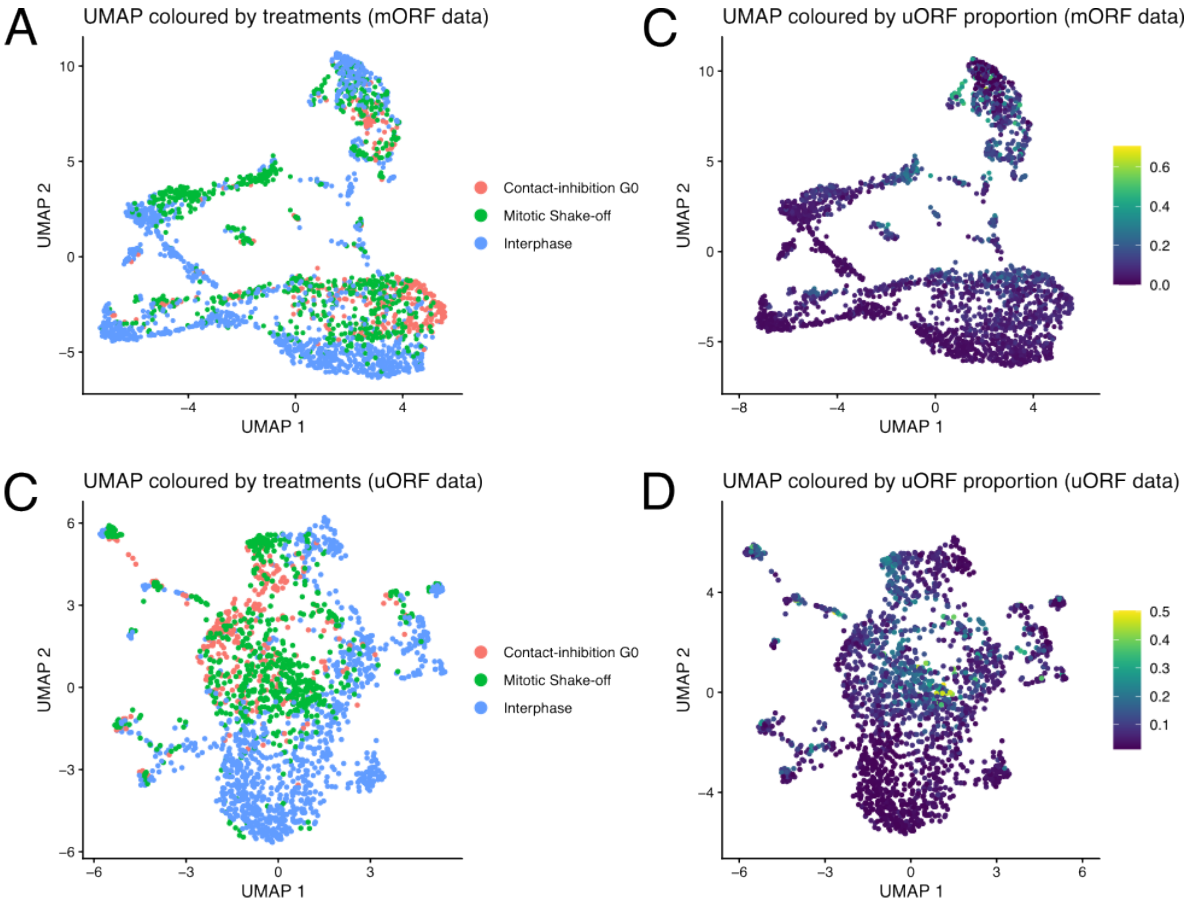


**Figure S1. Single-modality UMAPs (mORF-only and uORF-only) coloured by treatments and uORF proportion. (A)** UMAP derived from mORF-only features coloured by treatments. **(B)** The same mORF-only UMAP coloured by the per-cell uORF proportion $u/(u+m)$, where u and m are total uORF and mORF footprints per cell, respectively. **(C)** UMAP derived from uORF-only features, coloured by treatments. **(D)** The same uORF-only UMAP coloured by the per-cell uORF proportion $u/(u+m)$.
